# Wind drives temporal variation in pollinator visitation in a fragmented tropical forest

**DOI:** 10.1101/731273

**Authors:** James D. Crall, Julia Brokaw, Susan F. Gagliardi, Chase D. Mendenhall, Naomi E. Pierce, Stacey A. Combes

## Abstract

Wind is a critical factor in the ecology of pollinating insects such as bees. However, the role of wind in determining patterns of bee abundance and floral visitation rates across space and time is not well understood. Orchid bees are an important and diverse group of neotropical pollinators that harvest pollen, nectar and resin from plants. In addition, male orchid bees collect volatile scents that they store in special chambers in their hind legs, and for which the wind-based dispersal of odors may play a particularly crucial role. Here we take advantage of this specialized scent foraging behavior to study the effects of wind on orchid bee visitation at scent sources in a fragmented tropical forest ecosystem. We find that temporal changes in wind speed and turbulence are correlated with visitation to scent stations within sites, while local landscape structure is a strong determinant of spatial variation in visitation across nearby sites. These results suggest that the increased dispersal of attractive scents provided by wind and turbulence outweighs any biomechanical or energetic costs that might deter bees from foraging in these conditions. Overall, our results highlight the significance of wind in the ecology of these important pollinators in neotropical forests.

## Introduction

Animal pollinators such as bees provide critical ecosystem services that support biodiversity and global crop yields. Understanding the impacts of environmental change on bee communities is a central question for both the conservation of biodiversity [1] and agricultural productivity [2]. The composition and abundance of pollinator communities can vary substantially in both space [3] and time [4], likely as a result of both stochastic fluctuations and small-scale variation in the biotic and abiotic environment [5].

Wind strongly affects flying insects, and may be an important factor in spatio-temporal heterogeneity of pollinator visitation. Wind affects macroecological patterns of insect dispersal and migration [6,7]. Mean wind flow [8] and fluctuations (i.e., turbulence) pose biomechanical challenges [9–13] that push maneuverability limits in flying insects [14] and may impose energetic costs on flight [15]. The mechanical and physiological challenges posed by wind may have important effects on their interactions with plants, including herbivory [16,17], and pollinator visitation and landing [18]. Wind also indirectly impacts flying insects by inducing plant movements [19], which can impose additional maneuverability challenges [14]

In addition to biophysical challenges, wind also disperses chemical cues and signals critical for interactions between insects (e.g., attracting mates [20], or locating prey [21,22]), as well as between insects and plants (e.g., pollinator attraction [23–25] and herbivory [26]). Turbulence (i.e. fluctuations in wind on top of mean flow speed and direction) may be a critical factor for insects locating odor sources, as it disperses odors into complex plumes [27,28].

While wind is thus known to have many effects on pollinator behavior and ecology, our understanding of its influence on bee abundance and plant visitation rates is limited. While extreme wind speeds restrict bee flight and foraging (reviewed in [29]), recent studies have found that wind can have either positive [30] or negligible [10,31] effects on bee abundance and activity. Some wind’s variable effects could be explained by differences in pollinator sensory ecology, particularly the importance of olfactory cues for locating floral resources.

Orchid bees (Apidae:Euglossini) are a key group of pollinators for which wind-borne odors likely play a central role. The ~200 species (across four genera) of orchid bees are important neotropical pollinators of orchids and several other plant families [32]. In this group, foraging behavior intersects with mating strategies, as male orchid bees gather species-specific combinations of volatile compounds from a wide variety of flower species [33]. These fragrance “bouquets” are thought to play a key role in attracting mates [34]. Orchid bees are primarily forest-associated, and they collect fragrances from flowers and other sources that are often sparsely distributed within tropical rainforests. Accordingly, orchid bees show strong patterns of long-distance movement and dispersal across the landscape [35–38], and are thought to locate floral scent resources in dense vegetation using olfactory cues.

Despite its likely importance in dispersing scents, the role of wind in orchid bee ecology is not well understood, although previous observers have noted temporary increases in orchid bee arrivals at baits after wind gusts [39]. Recent work has also shown that male orchid bees performing mating displays strongly prefer to orient on the downwind side of trees, presumably to maximize the dispersal of odor plumes [40]. However, to our knowledge, quantitative studies of the impacts of wind (or turbulence) on spatial or temporal variation in orchid bee abundance and visitation rates at scent sources have not been made.

Wind may play a particularly important role in the fragmented forest landscapes that are increasingly characteristic of the Neotropical range of orchid bees. Previous work has found that forest fragment size can influence orchid bee abundance within fragments [41], despite the fact that bees move regularly between fragments. Forest fragmentation may affect orchid bees not only through direct impacts on habitat suitability, but also via indirect impacts on local wind patterns [42] and distribution of scent cues.

Here, we explore the effects of local landscape structure, wind speed, and turbulence on male orchid bee visitation rates to scent sources within a large tropical forest fragment and at adjacent deforested sites. In addition to a positive association between visitation and local forest cover, we predict that wind speed positively correlates with orchid bee visitation, because higher wind speeds will further disperse scents and attract bees from a wider area. Conversely, we hypothesize that turbulence is negatively correlated with visitation, because stronger turbulence will result in higher costs of flight and make odor plumes more challenging for bees to track.

## Methods

### Sampling sites and orchid bee collection

We collected male orchid bees from 9 different sites within or adjacent to a large forest fragment at the Las Cruces Biological Station (8.79°, -82.96°) in Coto Brus, Puntarenas Province, Costa Rica (Fig. 1). The forested areas in this region are characterized by premontane forest, with a high abundance and diversity of euglossine bees [41].

**Figure 1.**
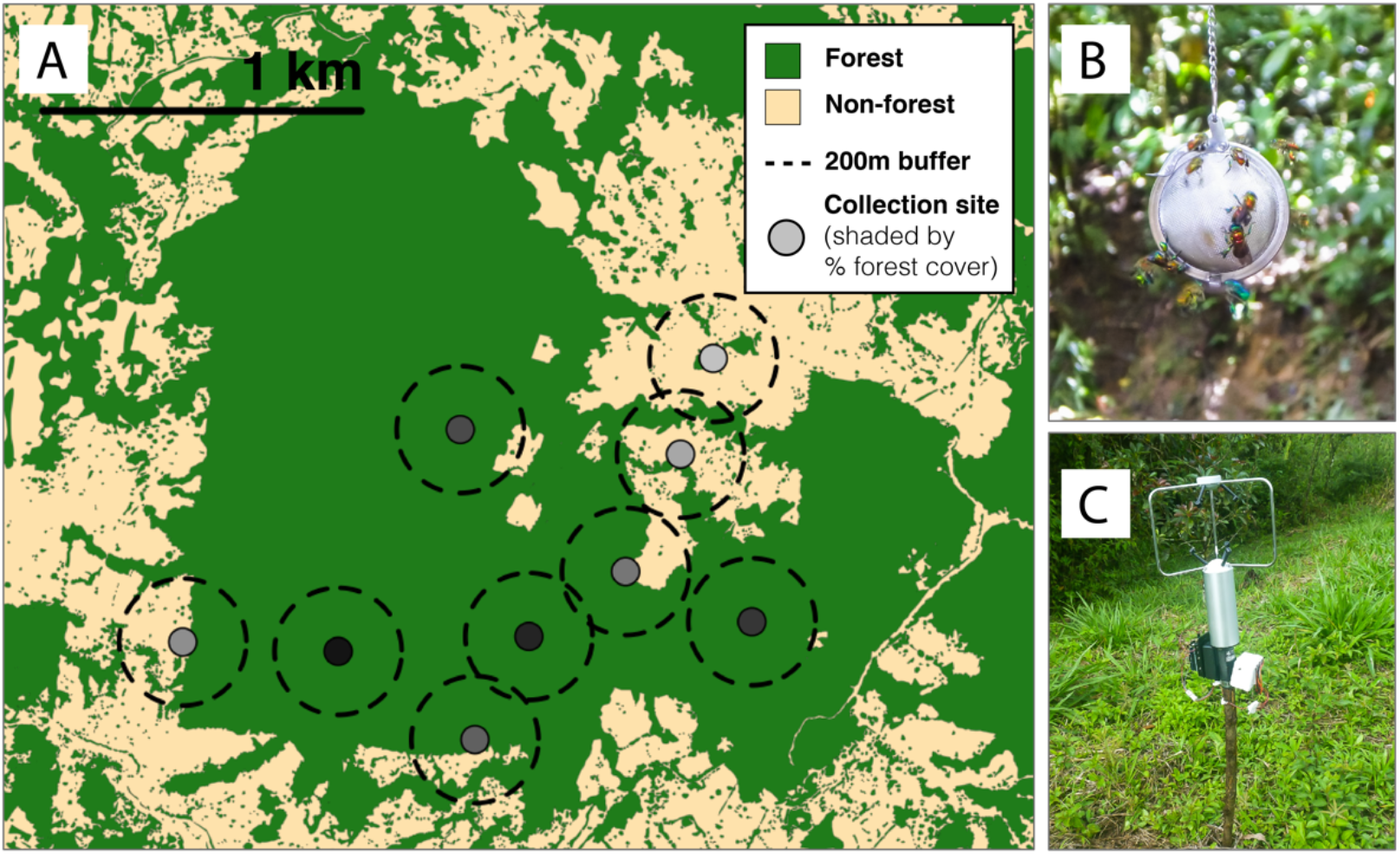
Study area and methods. (A) Map of collection sites (N = 9) at Las Cruces Biological Station, Puntarenas Province, Costa Rica. Solid markers show collection site (with shade indicating average percentage of local forest cover), and dotted lines show buffer zones used for measuring forest cover (radius = 200m). (B) Male orchid bees visiting a scent bait. (C) 3D Sonic anemometer deployed in the field. Photos: Julia Brokaw.

Male orchid bees were sampled from each site multiple (5-9) times between Oct 1 and Nov 16, 2014. All collections occurred between 8:30 and 11:30 a.m., roughly corresponding to peak daily abundance. Bees were sampled by saturating tissue paper with one of two compounds (cineole or methyl salicylate), and suspending this scent bait ~1.5 m above the ground in a permeable metal tea infuser. Male orchid bees arriving at the scent bait were collected by hand netting for 20 minutes. Bees (N = 409) were identified (Table S1) independently by two authors (JB and JDC) using an established taxonomic key [43]. Identifications of exemplar specimens were reviewed and corrected by Dr. Santiago Ramírez.

### Wind measurement

Simultaneously with each collection period, we characterized the local wind environment using a 3D sonic anemometer operating at 10 Hz placed 1 m above the ground and > 4m away from the scent bait. For each 20-minute wind sample, we calculated the mean wind speed and turbulence strength. We estimated the strength of turbulence by measuring the mass-specific turbulent kinetic energy of wind (*TKE*), as 0.5 * 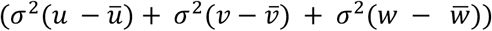, where *u, v*, and *w* represent wind speeds in the East-West, North-South, and vertical direction, respectively, and 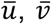, and 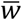, represent mean values along those respective axes. Overall mean wind speed was calculated as 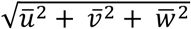. As turbulent kinetic energy correlated strongly with wind speed, we estimated the relative turbulent kinetic energy (hereafter “relative TKE”) as the residuals of a linear model of *TKE* on mean wind speed (log_10_(*TKE*)~log_10_(mean wind speed)).

### Estimation of landscape forest cover

To estimate the local forest cover at each site, we utilized a manually digitized GIS layer of smallscale (~2m resolution) forest elements in the region [44,45]. For each site, we calculated the extent of forest cover (% area forested) in concentric circles surrounding each site. We report data for a 200m buffer radius below, as this buffer radius was approximately nonoverlapping between sites, making landscape measurements largely independent across sites (Fig 1). Using radii ranging from 50-500m yielded qualitatively similar results (Fig S1).

### Data analysis and statistics

We built generalized linear models to test the relationship between forest cover and orchid bee visitation (median count values for each site) across sites. We built a generalized linear mixed model to test the effects of wind speed and turbulence (relative TKE) on orchid bee visitation. For mixed models, *p*-values were calculated using Satterthwaite’s approximation for degrees of freedom. Data and custom scripts are available on Zenodo (DOI: 10.5281/zenodo.3352365).

## Results

Orchid bee visitation varied significantly across sites (Kruskal-Wallis test, **χ**^2^ = 26.2, d.f. = 8, p = 9.7 x 10^-4^). We found no evidence that abundance differed between bait type (cineole vs methyl salicylate, generalized linear mixed model, N = 70, z = 1.017, p = 0.39), and collections from the two scents are combined below.

Orchid bee visitation was positively correlated with the percentage of forested area across sites (Figs 1, 2A, glm, df = 7, z = 1.97, p = 0.049).

**Figure 2.**
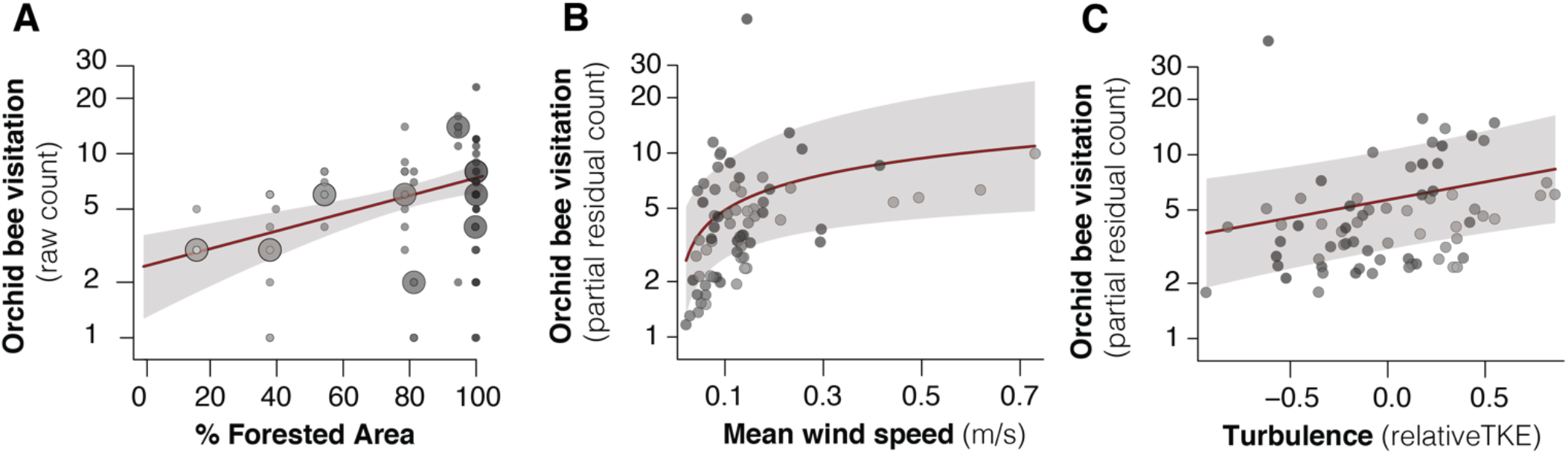
Forest cover and wind drive orchid bee visitation. (A) Effects of forest cover (% within 200m) on visitation across sites. Small and large markers show data for individual collections and site medians, respectively. Marker shade indicates amount of forested area (equivalent to site colors in Fig 1). (B-C) Marginal effect plots for the effects of wind speed (B) and turbulence (C) on orchid bee visitation. Solid markers show partial residuals for individual collections. In all panels, solid red line and shaded regions show the estimated relationship and +/− SE, from a generalized linear model (A), or generalized linear mixed model (B-C). Statistics for panels B-C reported in Table 1.

In addition, we found that both mean wind speed and turbulence had significant, positive effects on orchid bee visitation across time within sites (Fig 2B, Table 1). Across sites, however, wind speed was negatively associated with visitation (glmm, d.f. = 8, z = −1.96, p = 0.0495) and turbulence had no effect on visitation (glmm, d.f. = 8, z = 0.60, p = 0.55).

**Table 1.** Effects of wind speed and turbulence (relative mass-specific TKE) on orchid bee visitation across time within sites. Results are derived from a generalized linear mixed model with a poisson distribution, wind speed and relative turbulence as fixed effects, and site and bait scent (cineole vs methyl salicylate) as random effects.

| Variable | Estimated effect | z-value | p-value |
| --- | --- | --- | --- |
| Mean wind speed<br>($\log_{10}(\text{m s}^{-1})$) | 0.96 | 3.30 | **0.001** |
| Relative mass-specific TKE<br>($\log_{10}(\text{m}^2 \text{s}^{-2} \text{kg}^{-1})$) | 0.45 | 2.85 | **0.004** |

## Discussion

Our results suggest that both landscape and wind play important roles in driving male orchid bee visitation rates to scent baits in Neotropical forest habitats.

Forest cover was positively associated with orchid bee visitation across sites (Figs 1, 2A), consistent with the strong association between orchid bees and tropical forests [43]. Importantly, while previous work [41,46,47] has examined the role of fragment size in driving orchid bee abundance, our results show that local landscape structure within a large forest fragment also affect visitation rates. These results highlight the importance of detailed, small-scale landscape structure in driving biodiversity and ecosystem services, particularly in fragmented agricultural landscapes [44,45].

Our results also demonstrate that wind can influence orchid bee visitation over time within sites, despite the fact that measured wind speeds were quite low overall (median wind speed of 0.122 m/s). We found strong evidence that higher wind speeds are associated with higher visitation rates over time within sites (Fig 2B). The simplest explanation of this pattern is that increased wind speeds disperse attractive scents over greater distances, attracting male orchid bees from a wider area. Importantly, however, the positive association between wind speed and visitation within sites does not hold true between sites; visitation was weakly negatively associated with mean wind speed between sites. This likely results from forest-dominated sites (more typical of orchid bee habitat) having lower wind speeds on average.

We found no evidence that turbulence decreases orchid bee visitation; instead, higher relative turbulence was positively associated with visitation (Table 1). A possible explanation for this pattern is that turbulence also plays a role in dispersing attractive odors. This may be especially important when mean wind speeds are low, and scent dispersal occurs primarily through turbulent diffusion rather than bulk flow.

Overall, our results underscore the importance of wind for pollinator ecology, particularly in driving temporal variation in abundance and visitation rates of species that rely strongly on olfactory cues. Previous work has noted that orchid bee abundance can vary significantly across small spatial scales within the same habitat [48,49], as well as across time within sites [41]. Our results are consistent with these observations, and suggest that temporal variation within sites is driven, in part, by variation in wind speed and turbulence, consistent with previous anecdotal observations [46]. Future studies over a wider range of environmental conditions would help determine whether the positive relationship between wind and orchid bee visitation within sites changes with more extreme wind (e.g., in the presence of higher mean flows or strong gusts).

These results may also have important implications for the timing of volatile release in Euglossine bee-attracting plants. The release of pollinator-attracting fragrances from many flowers shows strong rhythmicity, likely synchronized with activity of pollinating animals [50]. Our results suggest that timing and efficacy of scent release by flowering plants could also be shaped by environmental wind conditions.

Understanding how wind interacts with landscape structure to drive pollinator abundance is critical for predicting the impacts of both local (e.g., forest fragmentation, [42,51]) and global (e.g., global shifts in wind and weather patterns [52]) environmental change on plant-pollinator communities. Our results suggest that changes in wind could have particularly important effects on temporal patterns of pollinator abundance and visitation rates at floral resources, which could have significant consequences for conservation and food production [53,54].

## Acknowledgments

The authors thank Randy Figueroa for help in orchid bee collection, Rodolfo Quiros, Zak Zahawi, and the staff at Las Cruces Biological Station for logistical and informational support, and Santiago Ramírez for help in species identification

This work was supported by a NSF GRFP fellowship to JDC, an Organization for Tropical Studies Pilot grant to JDC, a Company of Biology Traveling Fellowship to JDC, a David Rockefeller Center for Latin American Studies grant to JDC, NSF IOS-1257543 to NEP, and NSF CAREER IOS-1253677 to SAC.

