## Supplementary Material for "Wind drives temporal variation in pollinator visitation in a fragmented tropical forest"

### Supplementary Tables and Figures

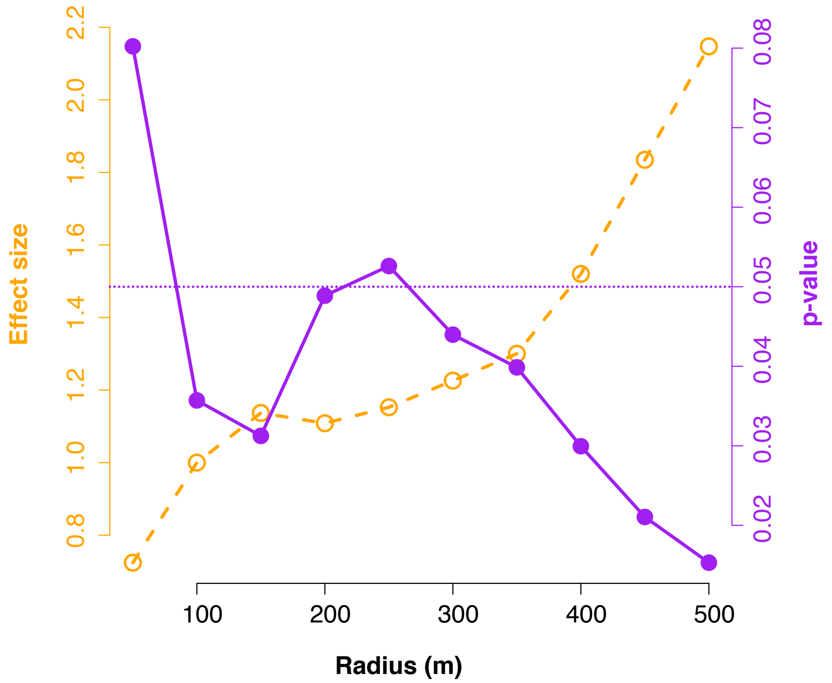


**Fig S1**. Sensitivity analysis for effects of forest cover buffer radius on orchid bee visitation. Estimated effect size (orange dashed line and open markers) and p-values (purple solid line and filled markers) from a generalized linear model ([median orchid bee count]~[% forest cover at a given radius]) for a range of test radii. Significance level of $p = 0.05$ shown as a dotted purple line.

**Table S1**. Species composition table. Species are listed in descending order of abundance across all collections.

| Species | # Cineole | # Methyl salicylate | Total | % |
|---|---|---|---|---|
| *Euglossa championi* | 115 | 7 | 122 | 27.1 |
| *Euglossa cybelia* | 70 | 0 | 70 | 15.6 |
| *Euglossa maculilabris* | 42 | 0 | 42 | 9.3 |
| *Euglossa flammea* | 39 | 0 | 39 | 8.7 |
| *Euglossa mixta* | 30 | 6 | 36 | 8 |
| *Euglossa dodsoni* | 30 | 0 | 30 | 6.7 |
| *Euglossa tridentata* | 18 | 0 | 18 | 4 |
| *Euglossa asarophora* | 17 | 0 | 17 | 3.8 |
| *Euglossa hansoni* | 7 | 7 | 14 | 3.1 |
| *Euglossa deceptrix* | 11 | 0 | 11 | 2.4 |
| *Euglossa gorgonensis* | 10 | 1 | 11 | 2.4 |
| *Euglossa sapphirina* | 8 | 0 | 8 | 1. 8 |
| *Euglossa bursigera* | 6 | 0 | 6 | 1.3 |
| *Eulaema speciosa* | 6 | 0 | 6 | 1.3 |
| *Euglossa erythrochlora* | 0 | 4 | 4 | 0. 9 |
| *Euglossa imperialis* | 3 | 0 | 3 | 0.7 |
| *Eulaema nigrita* | 3 | 0 | 3 | 0. 7 |
| *Euglossa despecta* | 2 | 0 | 2 | 0. 44 |
| *Euglossa heterosticta* | 2 | 0 | 2 | 0. 4 |
| *Euglossa purpurea* | 2 | 0 | 2 | 0. 4 |
| *Euglossa variabilis* | 2 | 0 | 2 | 0. 4 |
| *Euglossa allosticta* | 1 | 0 | 1 | 0. 2 |
| *Eulaema bombiformis* | 0 | 1 | 1 | 0. 2 |
